## Supplementary material for "*Plasmodium falciparum* serology: A comparison of two protein production methods for analysis of antibody responses by protein microarray": Supplmental Figure 1

**Supplementary figure 1.** Correlogram of multiple antigen-matched targets (left). Spearman's rank correlation reported ($r_s$) and increasing blue colour scale indicates relative strength of correlation based on calculated correlations for all proteins included in this analysis. Protein schematic (right) represents amino-acid aligned representation of IVTT (green) and purified (orange) proteins to the full-length native protein (grey). Proteins in the correlogram and schematic are correspondingly aligned.

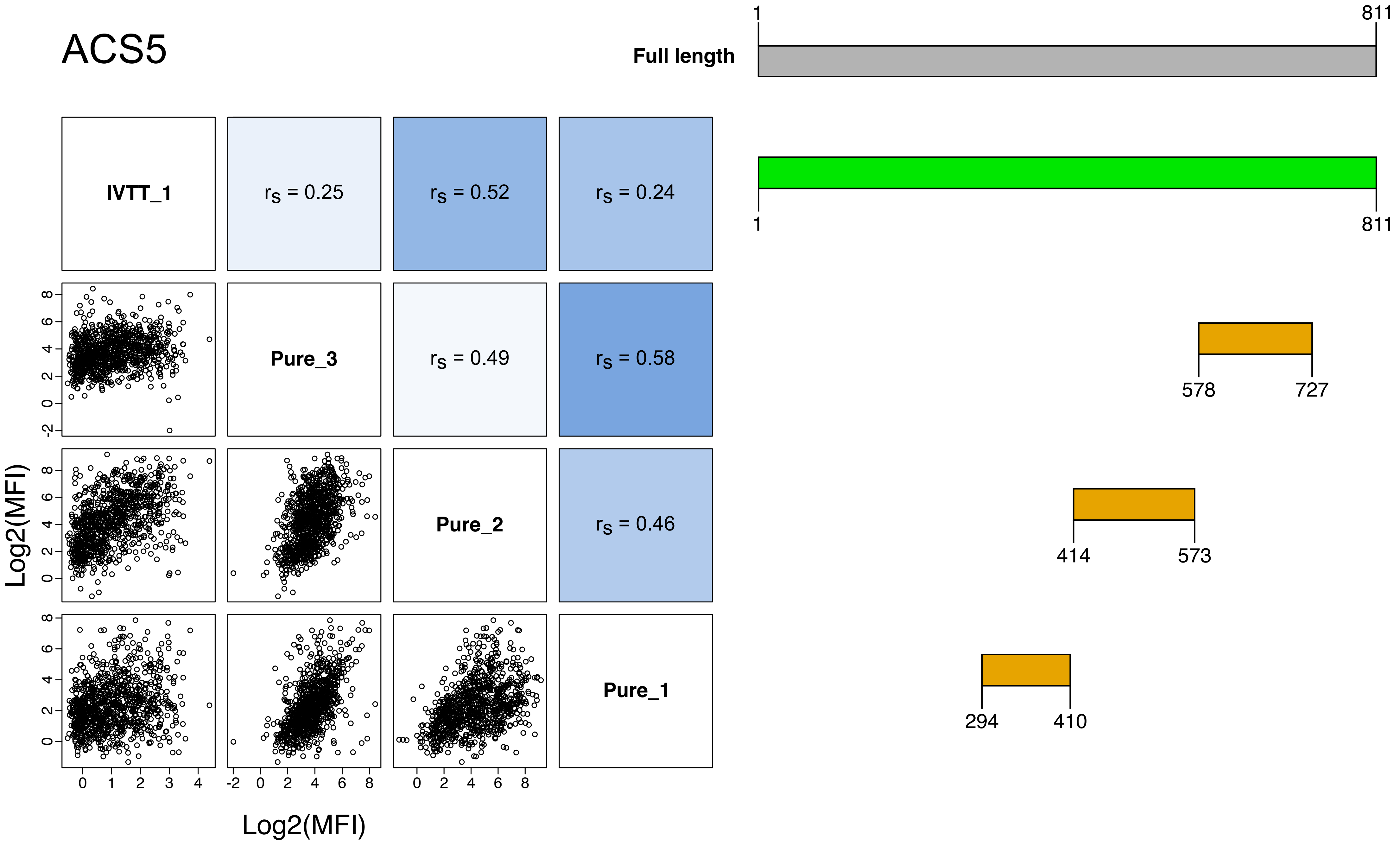

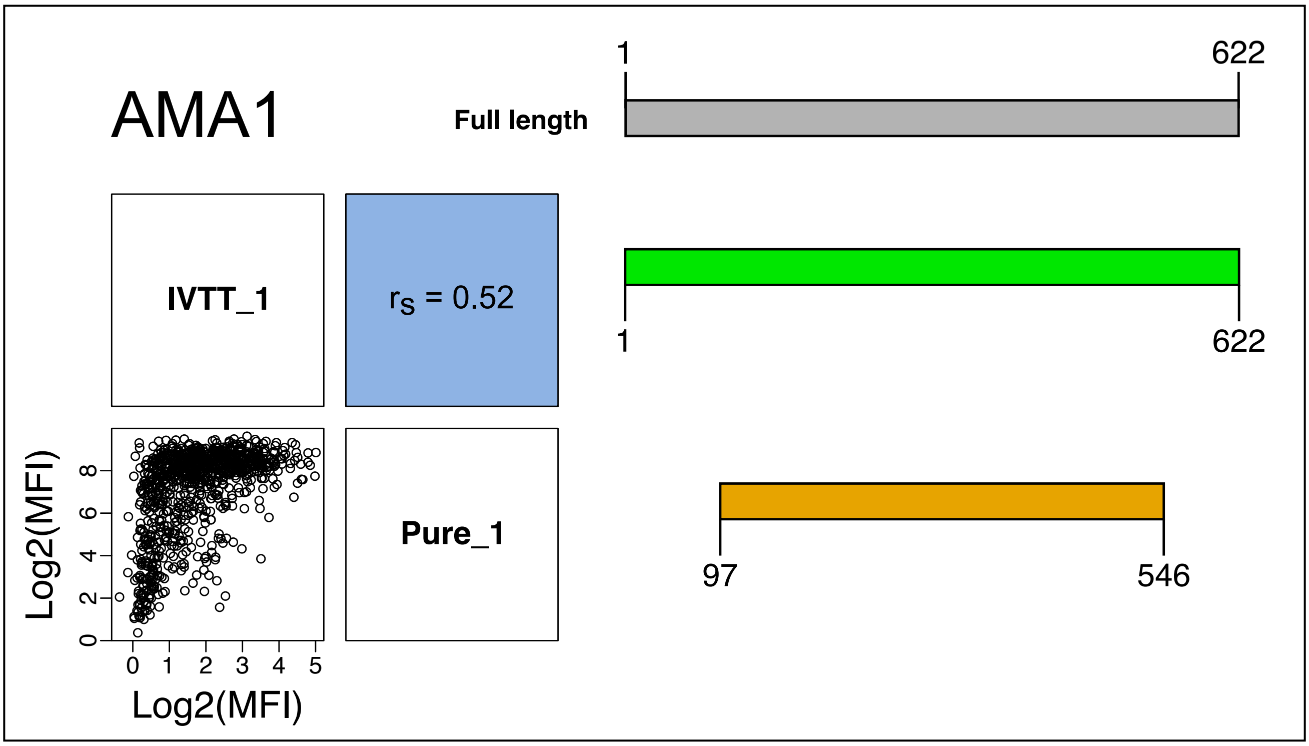
AMA1
Full length
1
622
IVTT_1
$r_S$ = 0.52
1
622
Pure_1
97
546
Log2(MFI)
Log2(MFI)

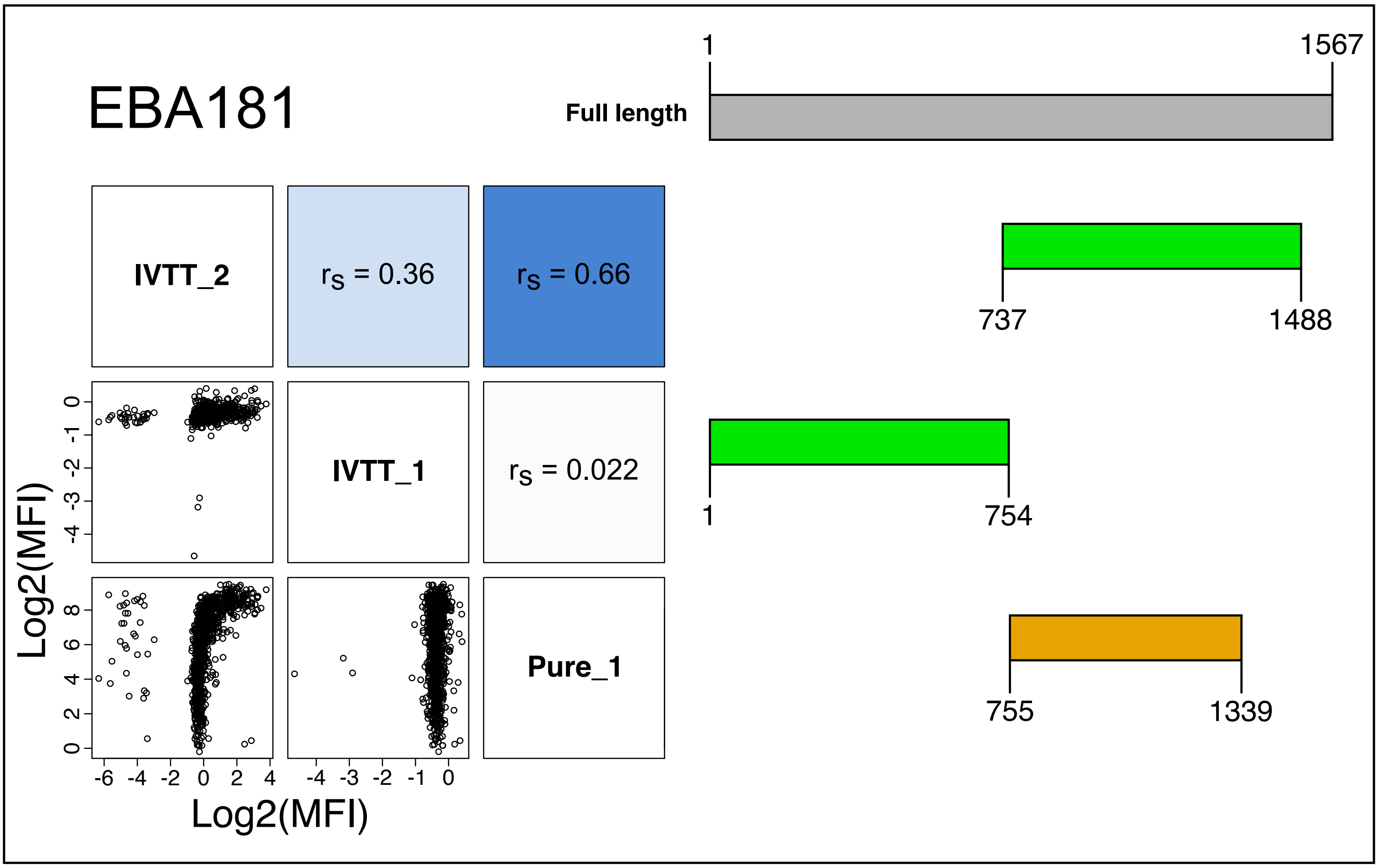
EBA181
Full length
1
1567
IVTT_2
$r_S$ = 0.36
$r_S$ = 0.66
737
1488
IVTT_1
$r_S$ = 0.022
1
754
Pure_1
755
1339
Log2(MFI)
Log2(MFI)

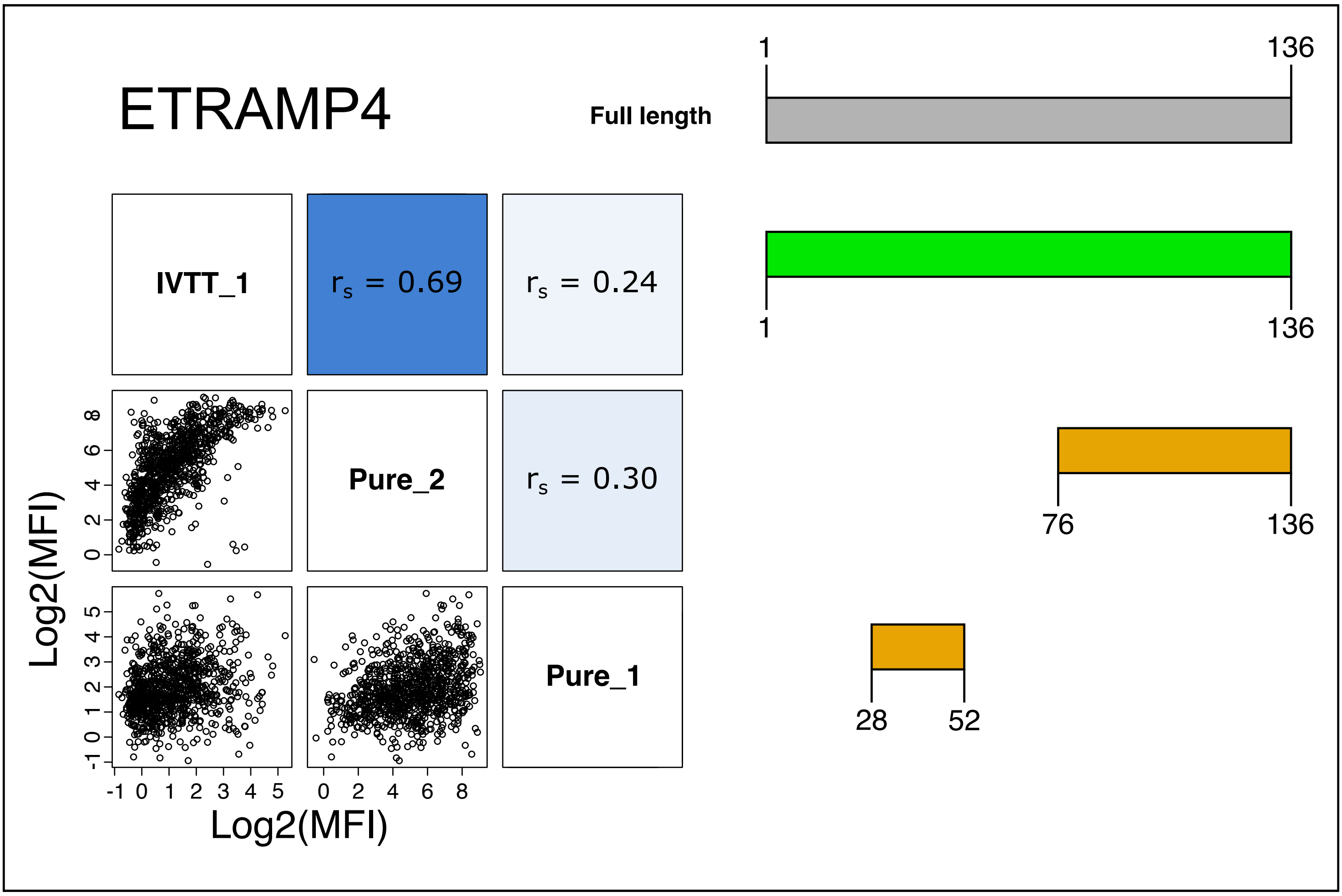
ETRAMP4
Full length
1
136
IVTT_1
$r_s$ = 0.69
$r_s$ = 0.24
1
136
Pure_2
$r_s$ = 0.30
76
136
Pure_1
28
52
Log2(MFI)
Log2(MFI)

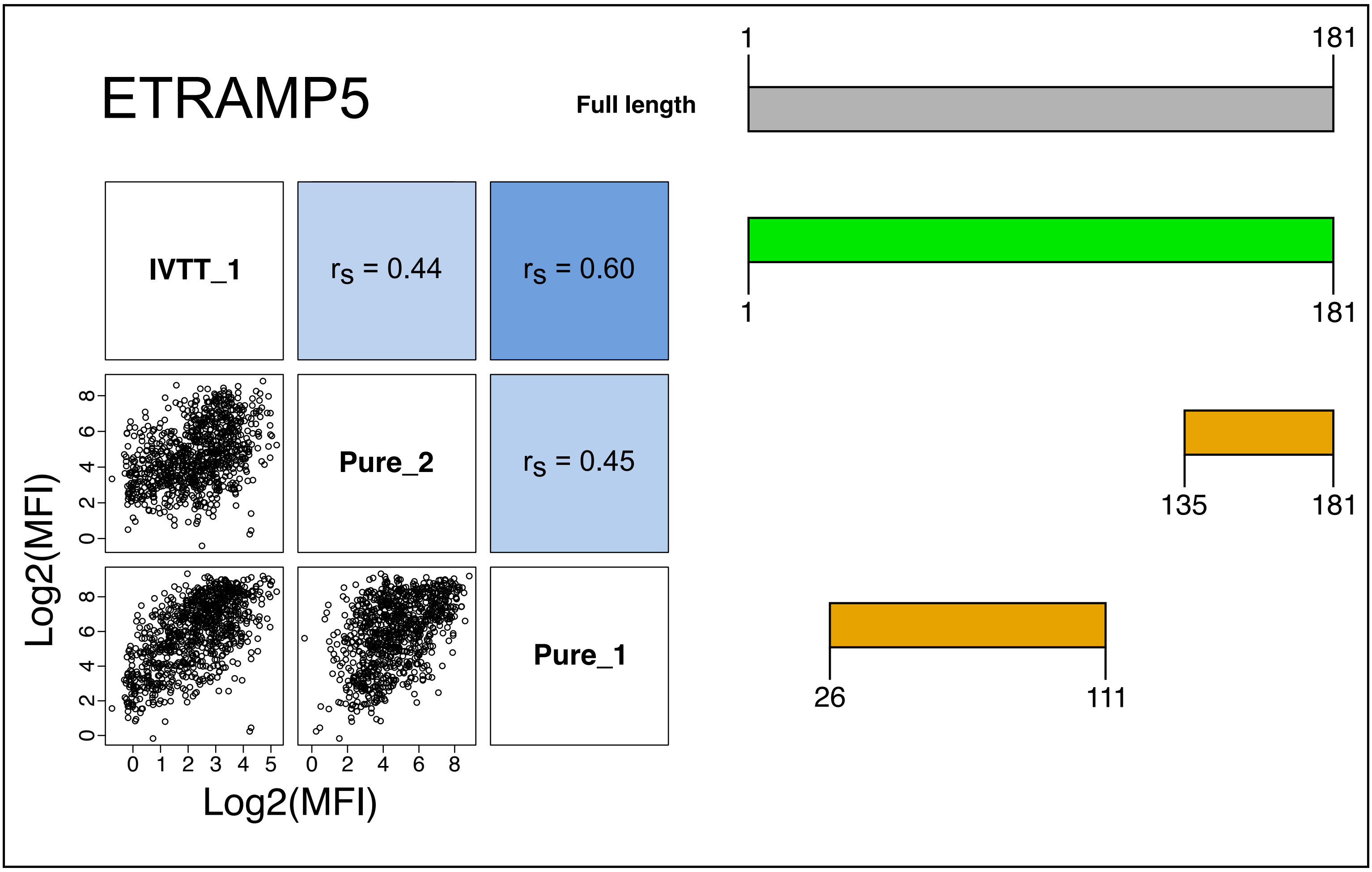
ETRAMP5
Full length
1
181
IVTT_1
$r_s$ = 0.44
$r_s$ = 0.60
1
181
Pure_2
$r_s$ = 0.45
135
181
Pure_1
26
111
Log2(MFI)
Log2(MFI)

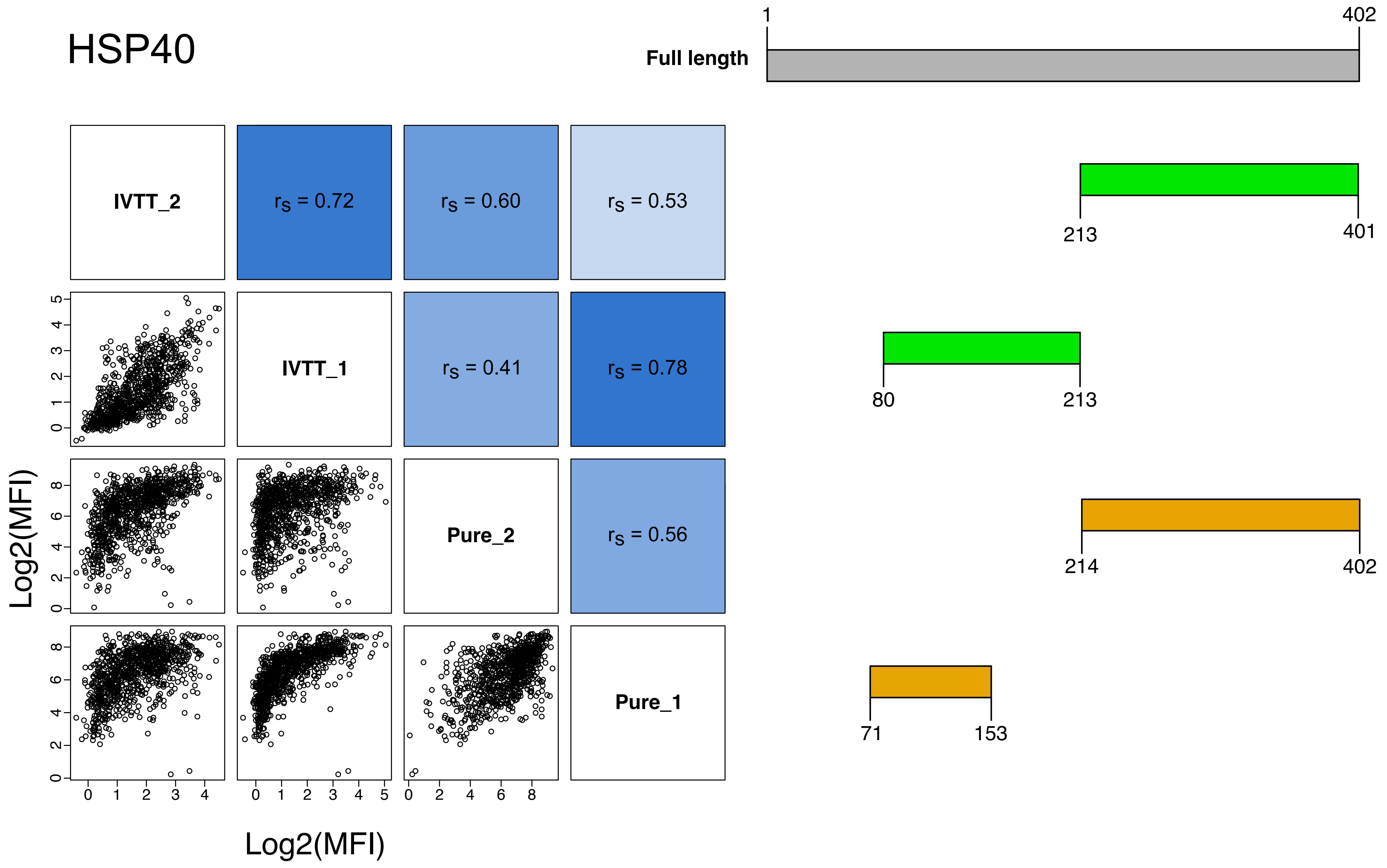

HSP40
Full length
1
402
IVTT_2
$r_S$ = 0.72
$r_S$ = 0.60
$r_S$ = 0.53
213
401
IVTT_1
$r_S$ = 0.41
$r_S$ = 0.78
80
213
Pure_2
$r_S$ = 0.56
214
402
Pure_1
71
153
Log2(MFI)
Log2(MFI)

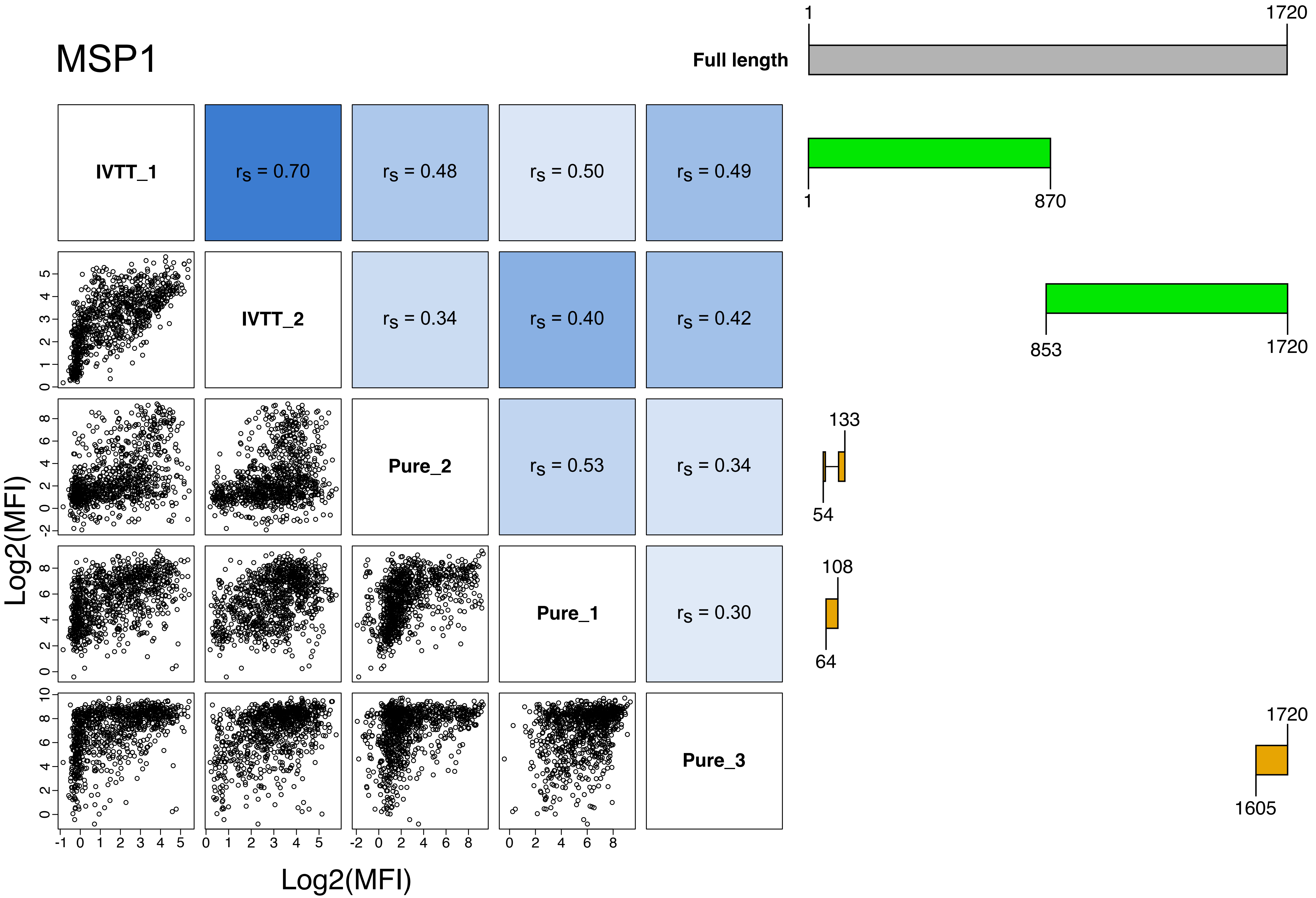
MSP1
Full length
1
1720
IVTT_1
rs = 0.70
rs = 0.48
rs = 0.50
rs = 0.49
1
870
IVTT_2
rs = 0.34
rs = 0.40
rs = 0.42
853
1720
Pure_2
rs = 0.53
rs = 0.34
133
54
Pure_1
rs = 0.30
108
64
Pure_3
1720
1605
Log2(MFI)
Log2(MFI)

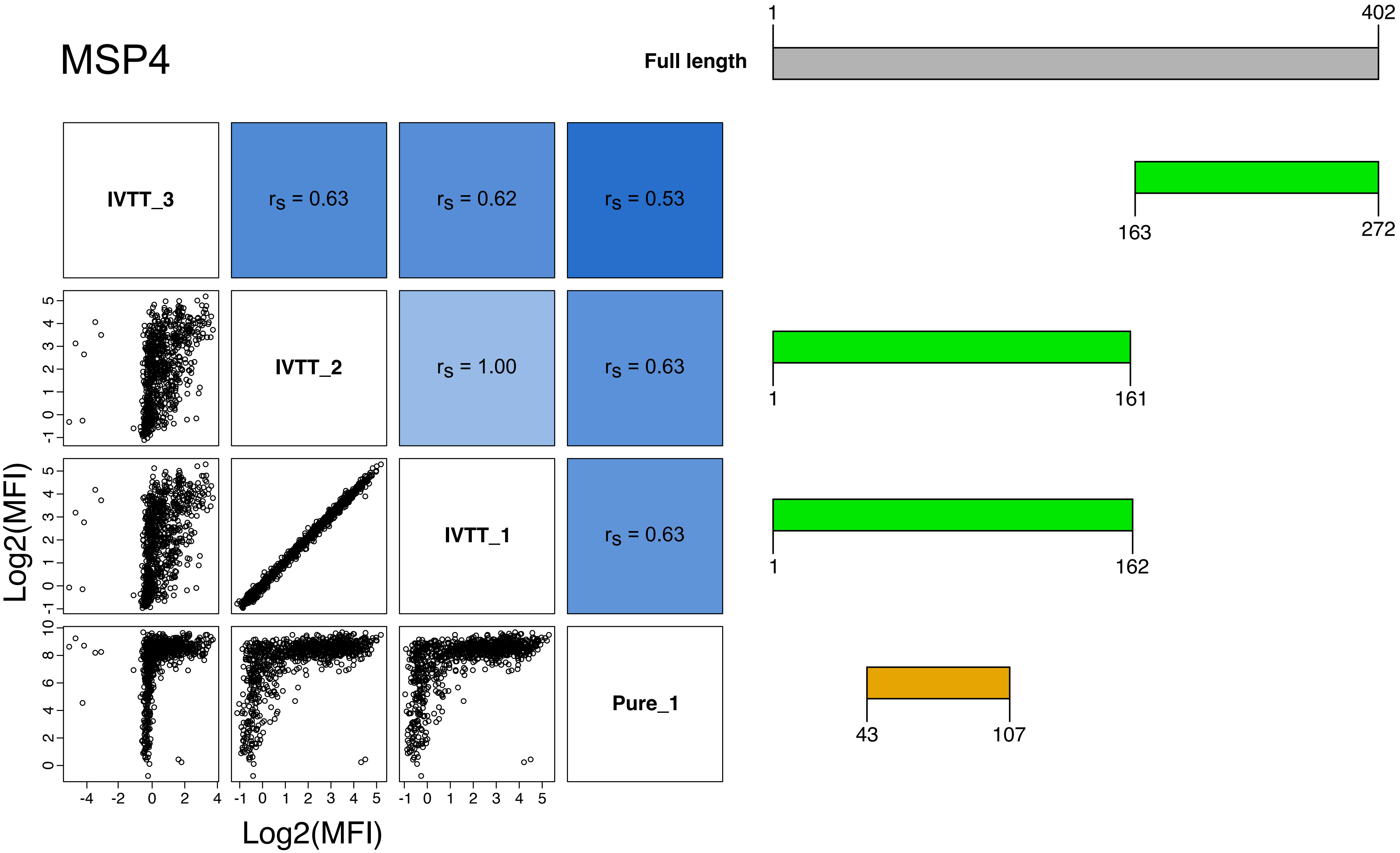
MSP4
Full length
1
402
IVTT_3
rS = 0.63
rS = 0.62
rS = 0.53
163
272
IVTT_2
rS = 1.00
rS = 0.63
1
161
IVTT_1
rS = 0.63
1
162
Pure_1
43
107
Log2(MFI)
Log2(MFI)

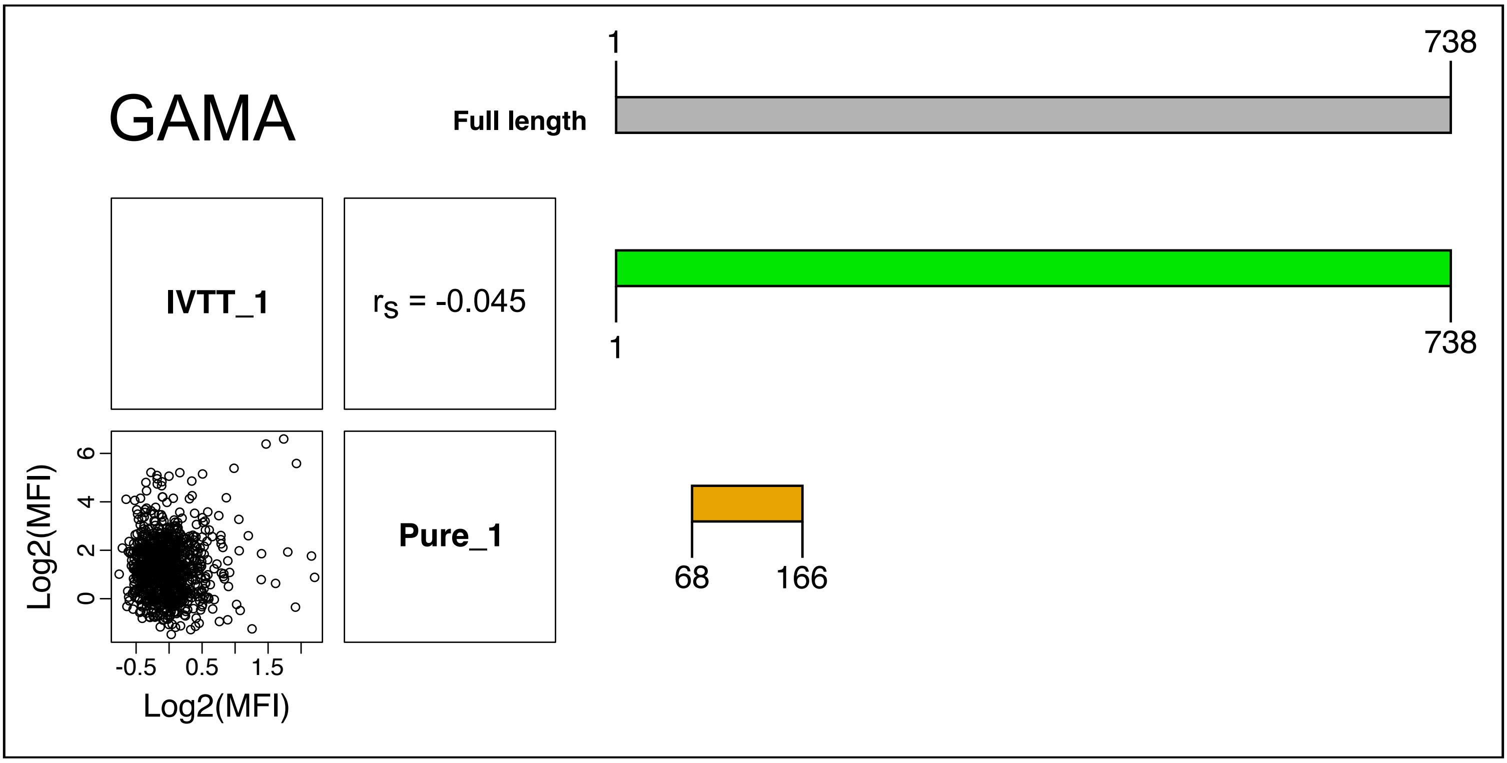
GAMA
Full length
1
738
IVTT_1
$r_S$ = -0.045
1
738
Log2(MFI)
6
4
2
0
-0.5
0.5
1.5
Log2(MFI)
Pure_1
68
166
