## Supplementary material for "*Plasmodium falciparum* serology: A comparison of two protein production methods for analysis of antibody responses by protein microarray": Supplmental Figure 2

**Supplementary figure 2.** Magnitude and range of response to IVTT and purified proteins, stratified by age. All sample responses (n = 899) to all protein targets grouped by antigen, presented with median and interquartile range.

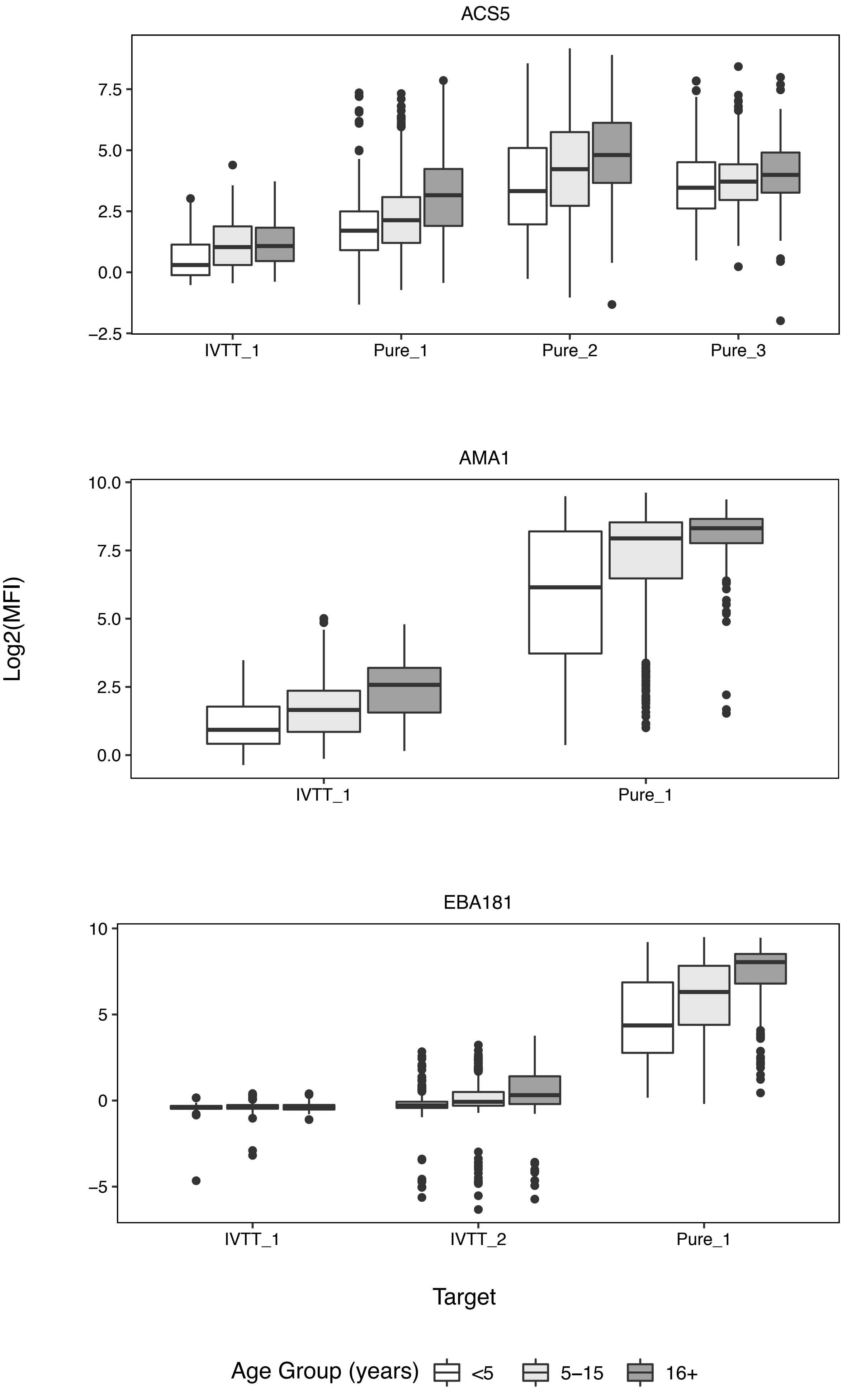

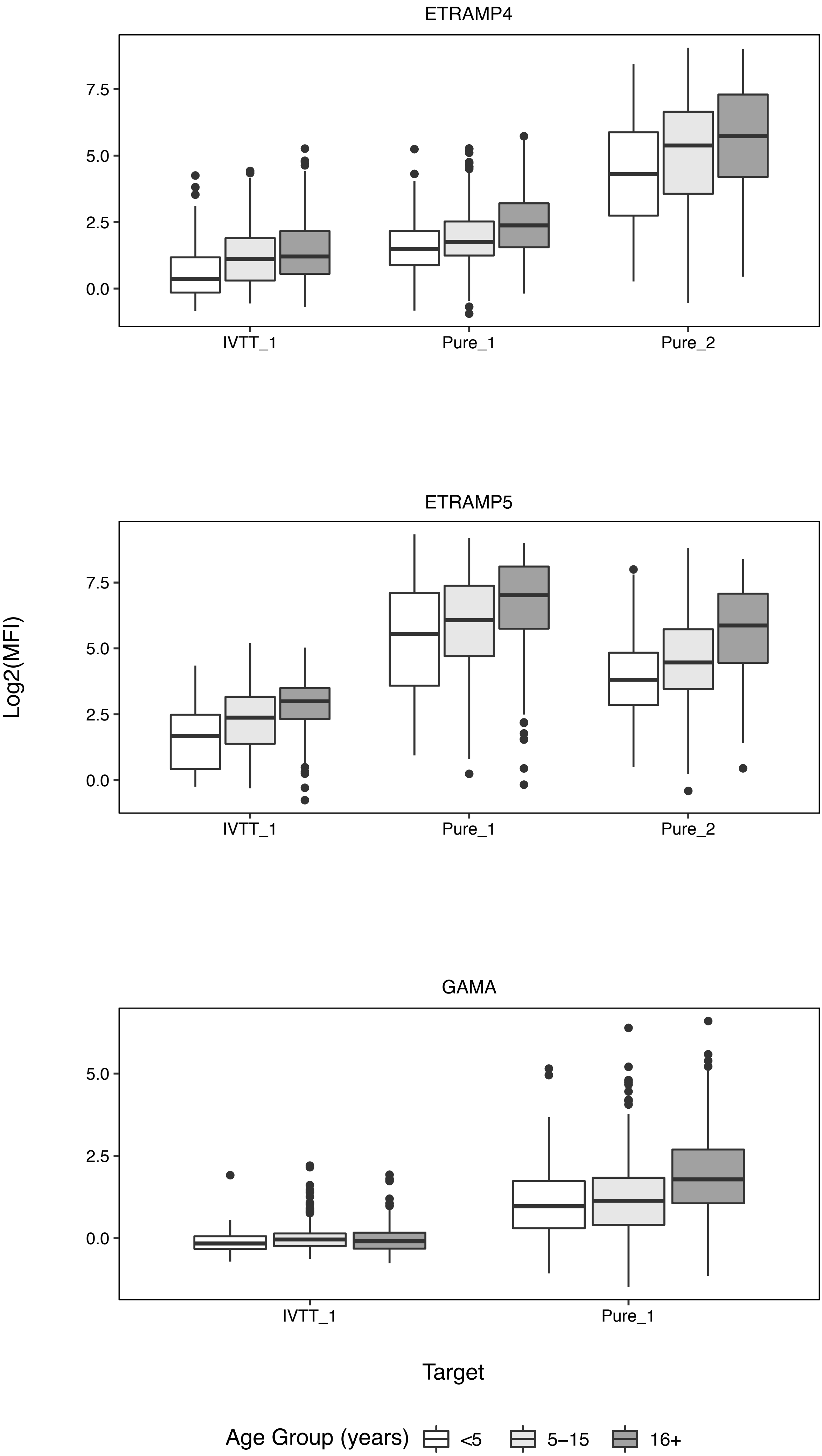

ETRAMP4
ETRAMP5
GAMA
7.5
5.0
2.5
0.0
IVTT_1
Pure_1
Pure_2
Log2(MFI)
Target
Age Group (years)
<5
5–15
16+

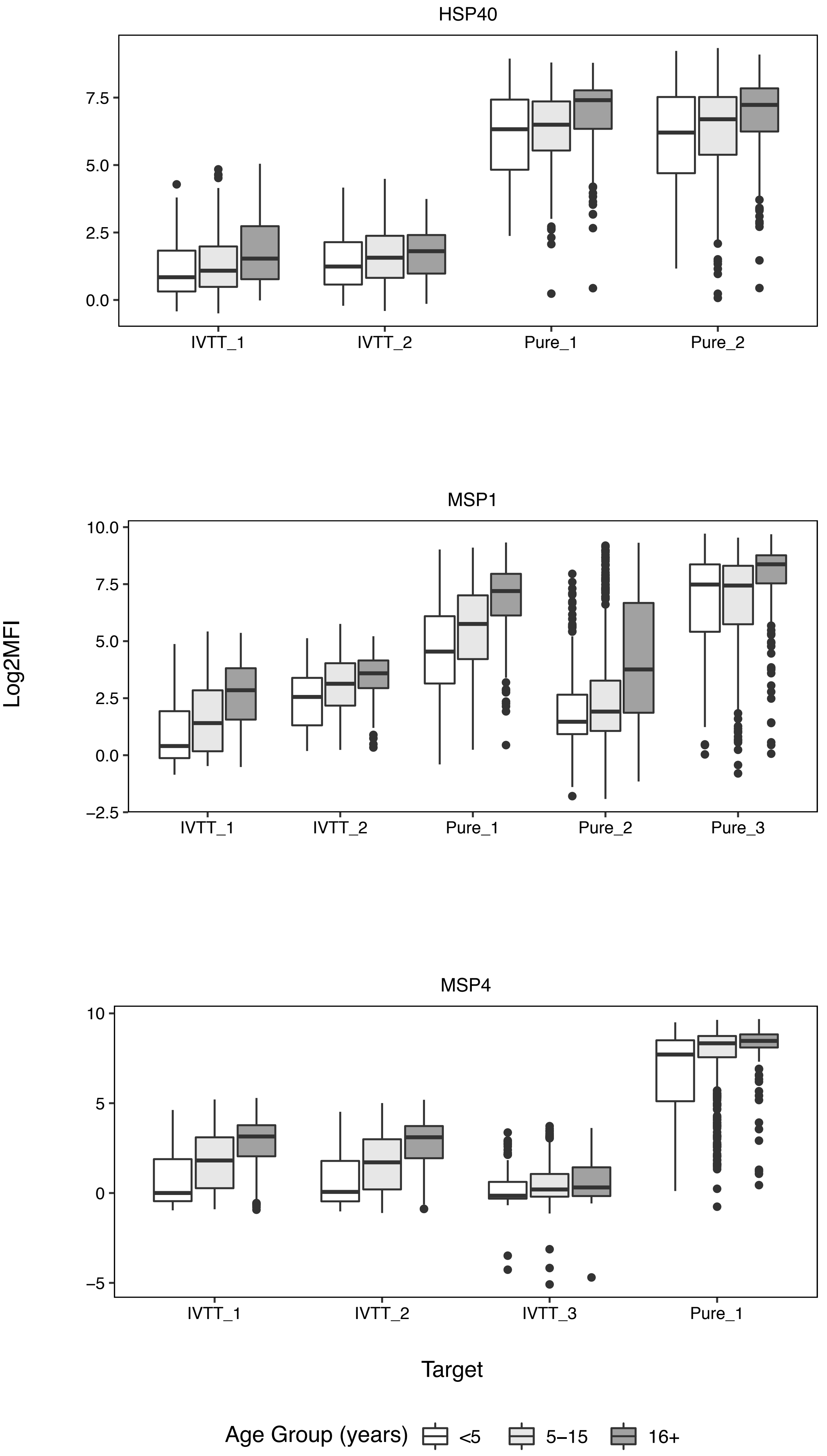

HSP40
7.5
5.0
2.5
0.0
IVTT_1
IVTT_2
Pure_1
Pure_2
MSP1
10.0
7.5
5.0
2.5
0.0
−2.5
IVTT_1
IVTT_2
Pure_1
Pure_2
Pure_3
Log2MFI
MSP4
10
5
0
−5
IVTT_1
IVTT_2
IVTT_3
Pure_1
Target
Age Group (years)
<5
5−15
16+

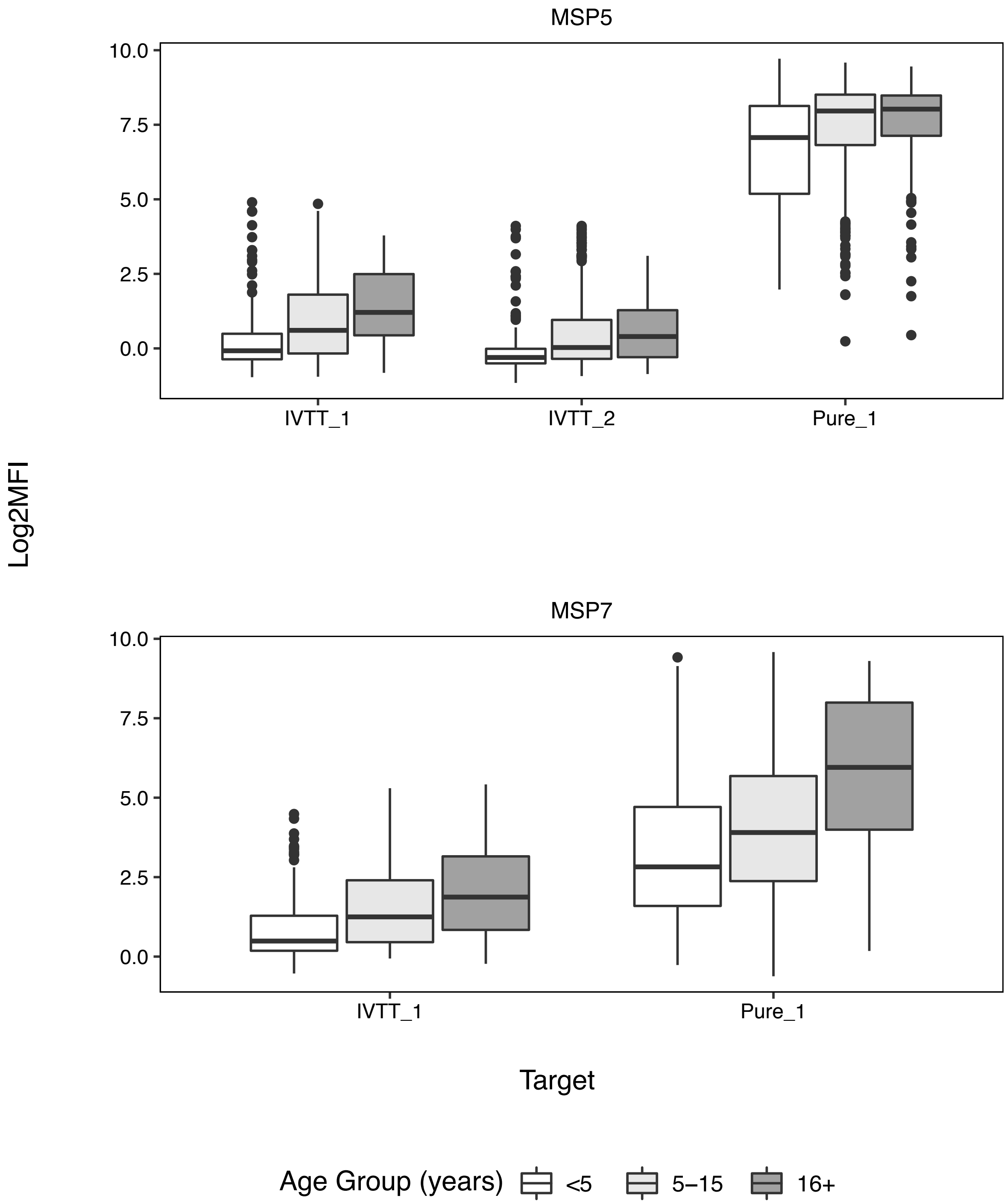
MSP5
10.0
7.5
5.0
2.5
0.0
IVTT_1
IVTT_2
Pure_1
Log2MFI
MSP7
10.0
7.5
5.0
2.5
0.0
IVTT_1
Pure_1
Target
Age Group (years)
<5
5–15
16+
