## Supplementary material for "*Plasmodium falciparum* serology: A comparison of two protein production methods for analysis of antibody responses by protein microarray": Supplmental Figure 3

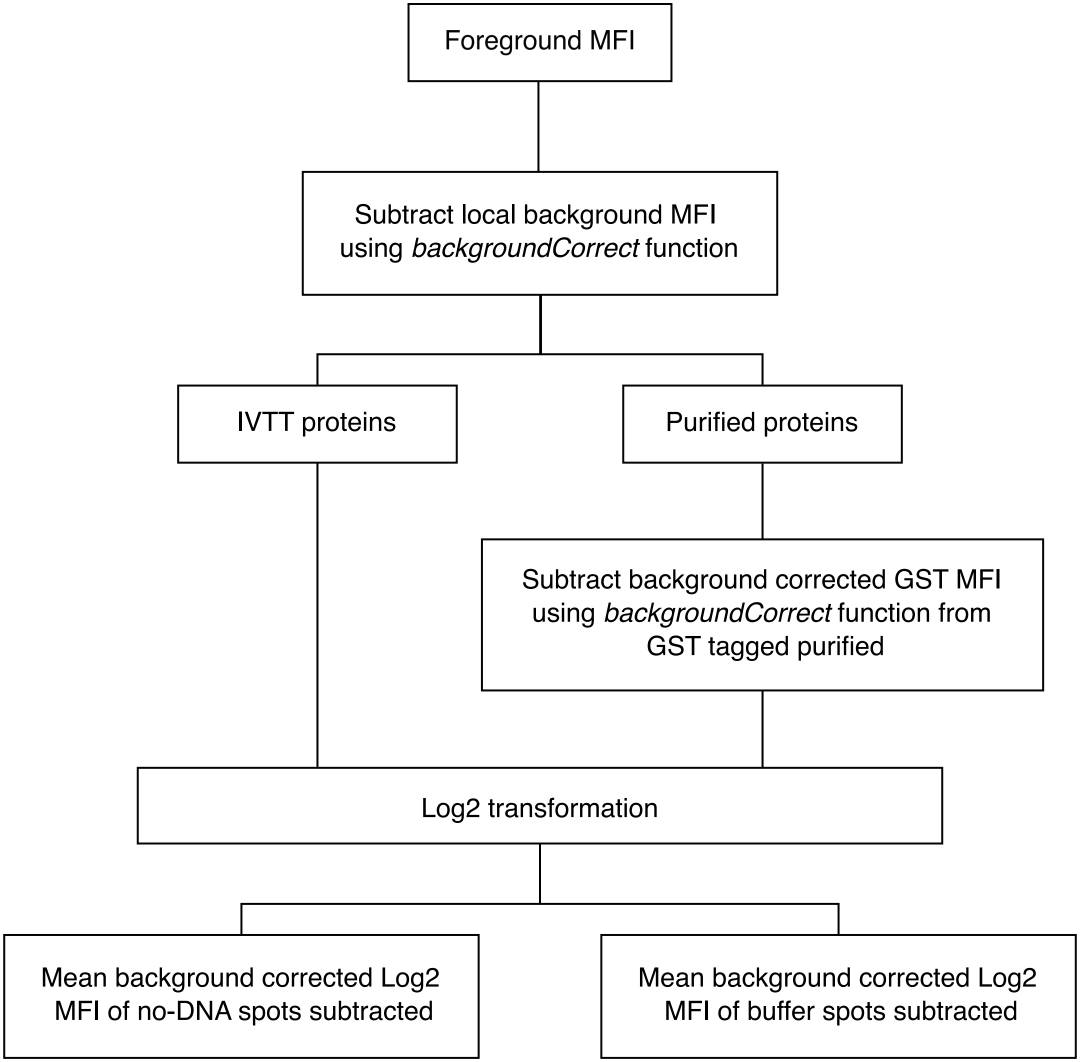


**Supplementary figure 3.** Data normalisation processes for IVTT and purified protein spots. After local background correction using the backgroundCorrect function from the limma package, purified protein spots were additionally corrected for possible GST reactivity by subtracting GST reactivity using the same function. After Log2 transformation, IVTT and purified proteins were normalised to background control spots of empty T7 vector and PBS buffer control spots respectively.
